# ChEBI-N - Enrichment Analysis with the ChEBI Ontology

**DOI:** 10.64898/2026.10.14.758784

**Authors:** Miranda Carlsson, Björn Gehrke, Charlotte Tumescheit, Adriano Rutz, Adnan Malik, Noel M. O’Boyle, Nicola Zamboni, Janna Hastings

## Abstract

**Introduction:** Biological and chemical ontologies are structured representations that can be used to map entities such as metabolites onto broader chemical and biological categories. Enrichment analysis applied to such ontologies identifies classes whose representation within a set deviates from an expected background distribution, helping to interpret complex metabolomics measurements.

**Objectives:** This study aimed to develop a tool for ChEBI (Chemical Entities of Biological Interest) ontology-based enrichment analysis, named ChEBI-N (ChEBI eNrichment). It is a successor of the previous tool BiNChE, with new features that broaden its applicability.

**Methods:** We reimplemented BiNChE and extended it to accept SMILES or InChI strings as well as ChEBI IDs. These broaden applicability to datasets with entities that are not yet present in ChEBI. We also constructed organism-specific background populations.

**Results:** We demonstrate the tool on three published metabolomics datasets covering wine, cheese, and human plasma samples. ChEBI-N correctly highlighted relevant enriched classes in all three datasets, such as flavonoids in the wine dataset and fatty acid classes in the cheese dataset. Comparing organism-specific and full-ChEBI backgrounds shows that restricted backgrounds reduce the inflated significance of generic classes, although the most enriched classes remain largely consistent.

**Conclusion:** ChEBI-N is an extended successor to BiNChE for ontology-based enrichment analysis of metabolomics data, offering a hypothesis-generating complement to existing metabolomics analysis pipelines.

**Availability:** ChEBI-N is freely available at chebin.hastingslab.org or as a Python package (pypi.org/project/chebin), with source code at github.com/ontology-tools/chebin, archived at Zenodo (doi:10.5281/zenodo.23264219) and Software Heritage (swh:1:rev: 133a41792704eb7f1db746f36a5ae97e9247b2b7).

## 1 Introduction

Metabolomics is the large-scale study of the full range of small molecules, or metabolites, in biological systems [1]. It has emerged as a powerful approach for characterising phenotypes and disease states [2]. Advanced analytical techniques, mainly Nuclear Magnetic Resonance (NMR) spectroscopy and Mass Spectrometry (MS), combine high sensitivity with broad coverage, making it possible to detect and quantify a wide range of metabolites and generating datasets of high complexity [1, 3]. This chemical diversity poses a significant challenge to biological interpretation; relating individual compounds to broader biological processes typically requires substantial manual curation and domain expertise. There is a need for computational tools able to map experimental results onto structured biological knowledge and support interpretation.

A bio-ontology is a structured representation of biologically relevant entities, linked through defined semantic relationships [4]. By encoding biological knowledge in a computationally accessible form, ontologies allow data, such as a list of genes or metabolites, to be systematically mapped onto the hierarchy [5]. This underlies ontology-based enrichment analysis, which assigns biological meaning to statistical patterns found in datasets by, for example, identifying entities that are differentially represented relative to a background dataset [6].

ChEBI (Chemical Entities of Biological Interest) is a well-established, manually curated and continuously updated ontology that organises knowledge on small molecules [7, 8]. This is done within three hierarchical structures: a molecular structure sub-ontology classified according to structural features, a role sub-ontology classified based on biological or chemical activity, and a subatomic particle sub-ontology which includes particles smaller than atoms [9]. At the leaf level, the structural hierarchy contains small molecules that may be measured in metabolomics experiments [6]. Within the hierarchy, classes are connected to broader classes by an *is a* relationship (rdfs:subClassOf) as Figure S1 shows. Molecules in the structural ontology can be connected to the role ontology through a *has role* relationship (RO:0000087); e.g. dopamine *has role* dopaminergic agent and *is a* catecholamine. Every class has its own ChEBI ID.

BiNChE [6] is a tool for ontology-based chemical enrichment analysis of small molecules, first introduced in 2015. BiNChE was based on the ChEBI ontology’s role and molecular structure hierarchies; the *subatomic particle* hierarchy was excluded [6]. It performed enrichment analysis to identify the statistically significantly overrepresented (enriched) classes in a user-defined set of molecules, relative to the background population of ChEBI classes and molecules [6]. Enriched classes were visualised as a graph-based network reflecting the structure of the ChEBI ontology with node colours corresponding to statistical significance. BiNChE was primarily developed to support the analysis and interpretation of metabolomics results [6]. In such studies, metabolites are experimentally obtained from biological samples; enrichment analysis on the identified metabolites based on ChEBI can provide a more comprehensive understanding of the underlying biology, by identifying which functional or chemical categories are overrepresented [6].

Ontology enrichment analysis is well established for the Gene Ontology (GO) [10], with tools such as BiNGO [11], topGO [12] and PANTHER [13]. For metabolomics, most available tools address other enrichments, including pathway enrichment analysis with MSEA [14] and MetaboAnalyst [15], which test metabolite sets against curated pathway definitions rather than against a structured chemical classification. The first tool for small-molecule enrichment analysis that uses ChEBI was MBRole [16], in which ChEBI is only used as one annotation source among others, with results presented as tables and summary plots rather than displayed on the ontology structure. To our knowledge, until now BiNChE [6] was the only tool that performed enrichment over the ChEBI hierarchy and returned an interactive graph-based visualisation.

BiNChE’s web interface is, however, no longer functional because its graph-rendering functionality depended on Adobe Flash Player, which was discontinued in December 2020. Furthermore, its outdated codebase is incompatible with modern software environments, limiting its continued use. These limitations, together with the opportunity to extend the tool with additional functionality, motivated the development of an updated version, ChEBI-N (ChEBI eNrichment). ChEBI-N is strongly based on BiNChE; it likewise uses the ChEBI ontology, but is implemented in Python (3.14) rather than Java, and introduces two key new features.

First, we implemented the possibility of using SMILES or InChI strings in addition to ChEBI IDs as inputs. Molecules that are not yet part of the ontology, and therefore lack a ChEBI ID, can be included through the integration of Chebifier [17], which predicts parent classes for molecules and includes them in the calculations. Broader input types may also simplify researchers’ workflows.

Second, ChEBI-N introduces the option to restrict the background to datasets narrower than the full ChEBI, with a *Homo sapiens*or *Arabidopsis thaliana*-relevant subset of the ontology implemented, as results may be more meaningful with a restricted background better suited to the specific study.

## 2 Implementation

Figure 1 provides an overview of ChEBI-N’s workflow, highlighting the steps performed by the user. In brief, the user provides a study-set of metabolites, and enrichment calculations are performed.

**Fig. 1.**
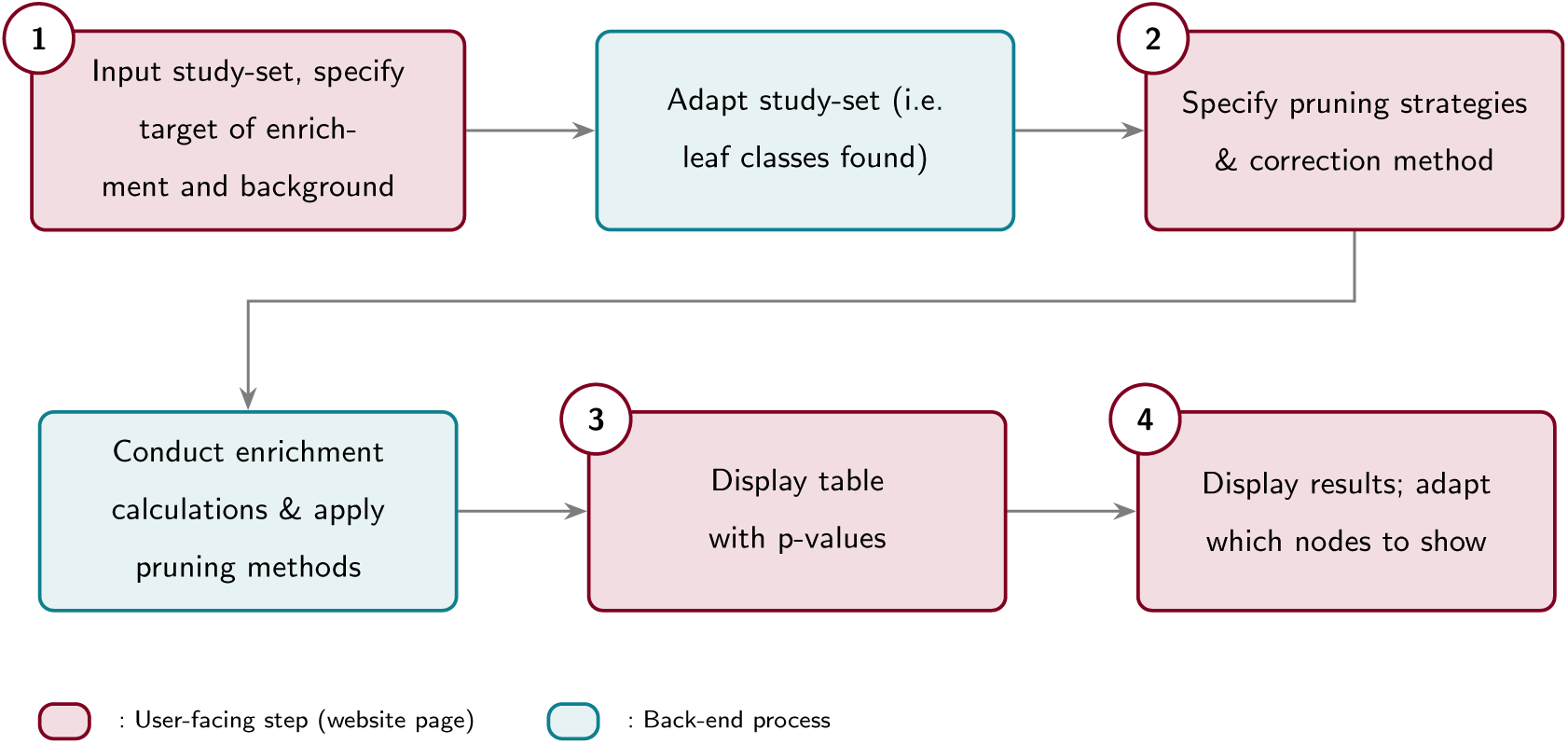
Workflow of ChEBI-N. Each number represents a new page on the website.

### Background population

The ChEBI ontology [7] is used as the default background, excluding any class flagged as deprecated. The ontology is downloaded in OWL format and loaded using the Py-Horned-OWL library [18]. Every month the underlying data is updated to the most recent version. As in BiNChE, calculations can be made on the molecular structure hierarchy (‘Structure’), the role hierarchy (‘Role’), or ‘Both’, referred to as ‘target of enrichment’. For the ‘Role’ target, all classes beneath the root node role (CHEBI:50906) are included. For the ‘Structure’ target, which is considerably larger, the root node chemical entity (CHEBI:24431) has four direct subclasses: atom, group, chemical substance, and molecular entity. Of these, we include only the branches of the latter two (CHEBI:59999 and CHEBI:23367, respectively), as the atom and group branches of the hierarchy are not expected to contain fully specified chemical structures.

In each of these cases, the background is constructed from the set of *structurally defined leaf classes*. In an ontology, a leaf is traditionally defined purely by its position: a class with no children. For enrichment analysis, however, what matters is whether a class corresponds to an actual molecule that could in principle be measured, so we adapt this definition. We call such classes *structurally defined*, and operationalise this as having a complete SMILES string: one that contains no asterisk (*). (In ChEBI, an asterisk marks an unspecified substituent or attachment point, so a SMILES containing one describes a broader chemical class rather than a single, fully specified molecule.)

We therefore adapt the hierarchy so that every structurally defined class is a leaf, regardless of whether it is a leaf in the original ChEBI hierarchy. Most structurally defined classes are already leaves, however, e.g. alanine (CHEBI:16449), with unspecified stereochemistry, has two children (L-alanine and D-alanine). alanine has a complete SMILES string and could itself be measured, so it should count as a leaf despite having children.

To achieve this, we flatten the hierarchy: whenever a structurally defined class has a structurally defined child, the edge between them is removed and the child is connected to the parent’s own parent instead. alanine, L-alanine and D-alanine thus become sibling leaves under the same parent class (Figure S2).

Using the full adapted ChEBI ontology as a background population is the standard option, but the possibility of using restricted, organism-specific backgrounds is also available. There are currently two *Homo sapiens*-based backgrounds, one broader than the other, and one *Arabidopsis thaliana*-based background. We chose *Arabidopsis thaliana* since it is the most studied model plant [19]. We built each organism-specific background from that organism’s metabolites. Concretely, each background consists of the leaf classes corresponding to those metabolites, together with all their ancestors.

The broader human background combined data downloaded from the Human Metabolome Database (HMDB) [20] at hmdb.ca consisting of metabolites found in the human body, with data queried from LOTUS [**?** ] at lotus-initiative.github.io consisting of chemical entities found in living organisms. LOTUS was filtered to *Homo sapiens* compounds and HMDB to ‘quantified’ and ‘detected’ compounds, excluding ‘expected’ ones. All compounds were mapped to ChEBI leaf classes, and those without a resolvable ChEBI ID were excluded. A ChEBI ID could be resolved through existing annotation, a SMILES lookup against ChEBI, an InChIKey match, or Chebifier-based classification (used for *∼*13% of the compounds). If a compound was not a leaf, it was replaces by its descendant leaf classes unless it had more than 150 leaf descendants, in which case it was excluded to prevent high-level classes from disproportionately inflating the background. When Chebifier provided several IDs, only the most specific (deepest) classes were kept. The resulting human background contains 7397 leaves from 4552 processed ChEBI IDs.

Although the data sources used to create this human background are based on compounds that have been found in humans, this does not mean the metabolites are endogenous. For research questions where an endogenous-based background may be relevant, we created a narrower *Homo sapiens*-based background from Recon3D [21], a resource from BiGG models [22]. Recon3D contains metabolites guaranteed to have the genetic machinery required to be synthesised in the human body. Compounds were mapped to ChEBI through their Recon3D annotations or, where none was given, through UniChem [23] via InChIKeys. This provided 2065 leaves from 797 resolved compounds.

The *Arabidopsis thaliana* background was constructed using the same workflow as the broader human background; however, only data from LOTUS was used, filtered by *Arabidopsis thaliana*. This produced 4253 leaves from 1460 filtered ChEBI IDs (where *∼*35% of the compounds required Chebifier classification).

An option when using a restricted background is to exclude any study-set leaf classes (the user input) that are not a part of the background. Since this may cause relevant classes to be dropped simply because the background happens to be incomplete, the default is to expand the background to include these automatically. Ideally, organism-specific backgrounds would already be comprehensive enough that this never happens, but given the data currently available, that is not the case.

### Input data

The study-set comprises the molecules of interest identified in a metabolomics study, provided by the user as either ChEBI IDs, SMILES or InChI strings (step 1 in Figure 1). If a ChEBI ID does not correspond to a leaf class, it is replaced by its descendant leaf classes. If a SMILES or InChI string cannot be matched to an existing ChEBI entry, its direct parent classes are predicted using Chebifier [17] and are resolved to leaf classes in the same way as for the backgrounds. Weights, indicating for example the relative importance of each compound, may be added to the study-set, changing how the enrichment calculations are conducted. If a non-leaf class is assigned a weight in the input, the corresponding leaves each receive the same weight as that parent.

### Enrichment analysis calculations

All classes in the chosen hierarchy and background that are ancestors of the classes in the study-set receive a p-value, except those removed by pruning. For an unweighted study-set, Fisher’s exact test is used for p-value calculations as described in Supplementary Section S4.

### Pruning strategies

Pruning options, i.e. filtering of nodes, are available mainly to make the final graph less cluttered (step 2 in Figure 1). Pruning methods can either be used once each or combined in a predefined loop. The pruning methods are explained in Supplementary Section S5.

### Correction method

Multiple hypothesis testing correction is used to reduce the number of false positives, since the likelihood of these errors increases as the number of statistical tests increases [24]. The options available in ChEBI-N are Benjamini-Hochberg [25], Bonferroni [26], and none, of which Benjamini-Hochberg is recommended.

### User interface

The web framework is implemented using Flask. Results can be downloaded as a CSV file, and are shown as a table of raw and corrected p-values from which an interactive graph view can be generated. Graphs are rendered with Cytoscape.js [27]; darker nodes are more significant on either a relative (default) or absolute *−* log_10_(*p*) scale. Corrected p-values are used where available. The layout can be changed, and the graph exported as PNG, JPEG, or PDF. Hovering over a node shows its ChEBI ID and raw and corrected p-values; nodes can be selected individually or together with their neighbours or descendants, and moved by dragging (Figure S3).

## 3 Results and discussion

ChEBI-N is available as a web interface at chebin.hastingslab.org, and as an installable package (pypi.org/project/chebin). The code is available at github.com/ontology-tools/chebin (v1.1.0: doi:10.5281/zenodo.23264219, swh:1:rev:133a417).

We applied ChEBI-N to data from three studies with varying use cases found on MetaboLights, an open-access database for metabolomics studies [28]. All studies are identified by a MetaboLights ID: MTBLSxx. The study-sets have been chosen to be of reasonable size to simplify interpretability and to contain entities that many may recognise (wine- and cheese-based), as well as a human dataset. An evaluation of ChEBI-N’s performance on a larger dataset (564 metabolites) on ribosomal mistranslation in humans [29] is available in Supplementary Section S7. All provided enrichment analyses are based on data accessed on 01.10.2026.

### Wine dataset from Arapitsas et al. [30]

The study MTBLS55 [30] compares wine stored domestically and in a cellar across 54 annotated metabolites, given as both ChEBI IDs and SMILES. Figures 2 and S4 show the ChEBI-N enrichment graphs for this dataset (ChEBI IDs) based on the structure hierarchy; Figure 2 applies the plain enrichment pruning strategy, while Figure S4 uses default settings with no pruning applied. Iterative pruning results in a substantially less cluttered graph while retaining the most statistically and biologically relevant nodes. The following discussion focuses on the pruned results.

**Fig. 2.**
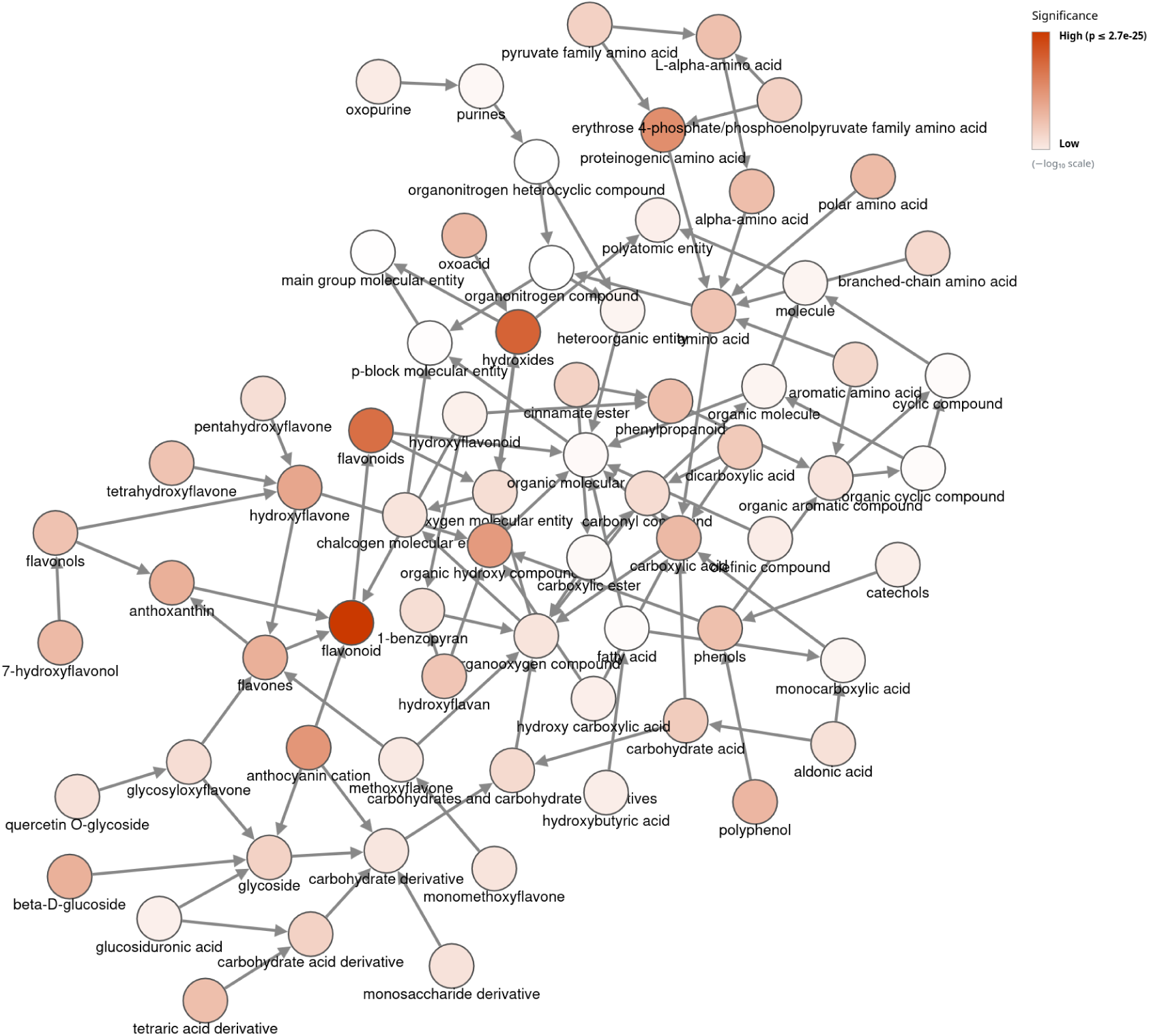
ChEBI-N enrichment analysis graph of the study-set from MTBLS55: *The influence of storage on the “chemical age” of red wines* [30] using ChEBI IDs. The results are only on the ‘Structure’ hierarchy, and the plain enrichment pruning strategy has been applied.

Key changes in domestically stored wine included formation of monosulfonated flavanols, accelerated transformation of grape-derived anthocyanin pigments, and an increase in pinotin A-like pigments, alongside a decrease in colour due to anthocyanin degradation [30]. The ChEBIN enrichment results reflect the compound classes underlying several of these findings. The most significantly enriched class, indicated by its dark red colour in the graph, is flavonoid (CHEBI:47916, *p_corr_* = 2.65 *×* 10*^−^*^25^), consistent with the dominance of anthocyanins, flavanols, and flavonols as primary markers in the study [30]. Within this, anthocyanin cation (CHEBI:35218, *p_corr_* = 8.81 *×* 10*^−^*^14^) reflects the central role of grape pigments, reported to be significantly higher under cellar conditions [30]. Several amino acid classes are also enriched, including proteinogenic amino acid (CHEBI:83813, *p_corr_* = 9.35*×*10*^−^*^15^) which is explained by the chemical composition of wine rather than a storage-dependent biological signal.

Next, we compared the results of using ChEBI IDs vs SMILES as an input when applying the same pruning strategy (plain enrichment pruning). All SMILES could be matched to an existing ChEBI class, so no Chebifier predictions were needed. We found 61 classes common to both analyses, nine unique to the ChEBI ID input and six unique to the SMILES input. For 58 of the 61 shared classes, the raw p-values were identical. The most enriched class remained flavonoid in both cases, with *p_corr_* = 2.54 *×* 10*^−^*^25^ for the SMILES input and *p_corr_* = 2.65 *×* 10*^−^*^25^ for the ChEBI ID input. Differences between ChEBI ID and SMILES inputs can be explained by some SMILES not matching the same ChEBI IDs as intended. When a SMILES matches multiple ChEBI IDs, only one is included in the analysis (always the lowest ID, for the results to stay reproducible). This occurred for five of the SMILES, and in one case a different ID than in the ChEBI ID input was selected. Furthermore, some SMILES do not have the stereochemistry information that their intended ChEBI ID specifies, causing them to match a different class. This explains the three shared classes with different p-values, all of which are amino acid classes.

The results show that using SMILES as input is a viable option, but it may be important to verify that matches are as specific as intended, and manual lookup of certain ChEBI IDs may sometimes be more reliable. Since ChEBI-N allows a mixed input of SMILES and ChEBI IDs, users can for example selectively double-check SMILES with multiple matches as these are reported alongside the enrichment results.

### Cheese dataset from Caron et al. [31]

To further explore the usage of ChEBI-N, we analysed the metabolites measured in the study MTBLS1509 [31]. The study compares cheeses produced with strains from four genetically differentiated *P. roqueforti* populations [31]. We used the 63 metabolites reported in the study, each annotated with a ChEBI ID, as the study-set for both structure and role-based enrichment on the full ChEBI background. The resulting enrichment graph is shown in Figure 3. We applied three levels of root-children pruning, high p-value pruning with a threshold of 0.05, and zero-degree pruning, and used Benjamini-Hochberg p-value correction.

**Fig. 3.**
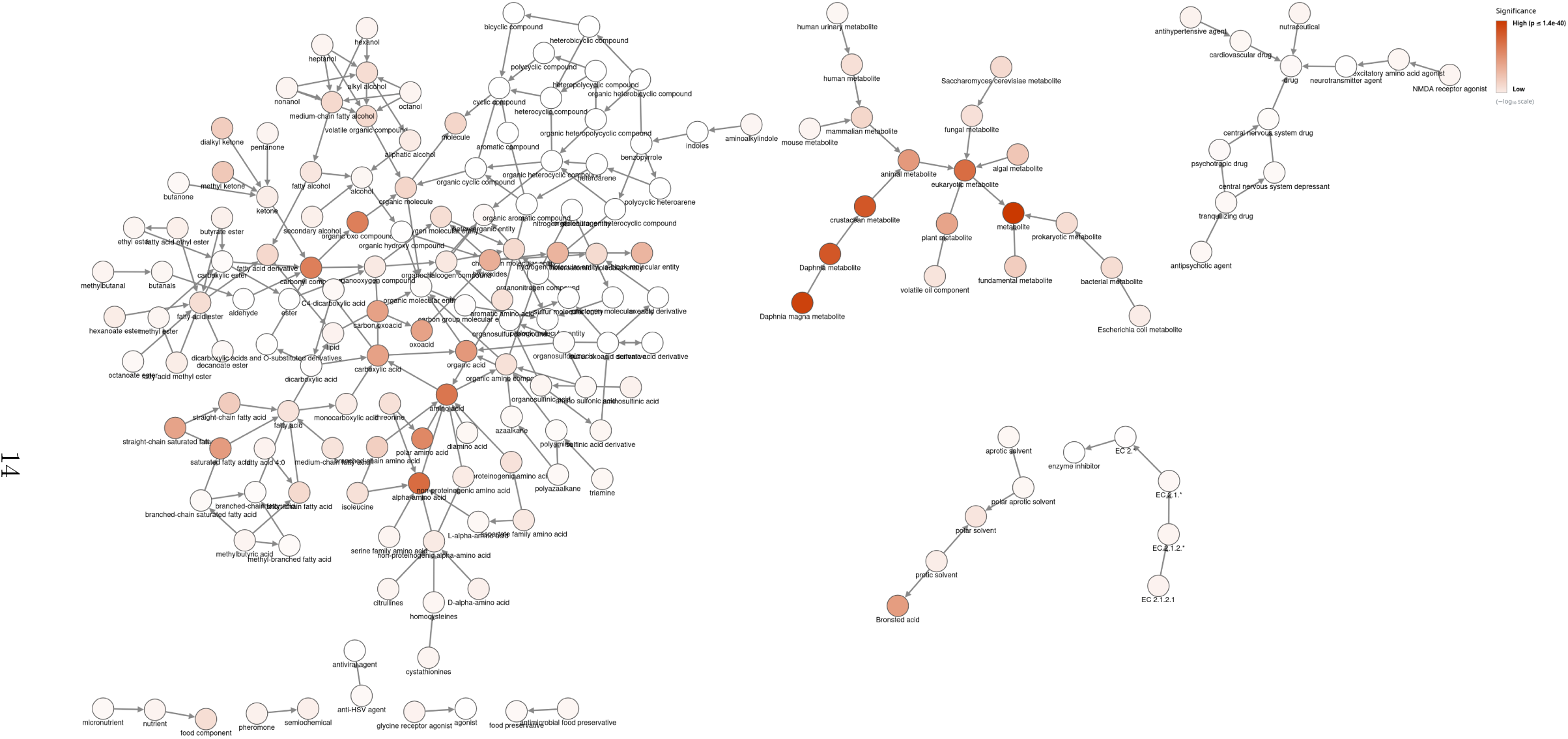
Enrichment analysis of the study-set from MTBLS1509: *Strong effect of Penicillium roqueforti populations on volatile and metabolic compounds responsible for aromas, flavour and texture in blue cheeses* [31] on both ‘Role’ and ‘Structure’ hierarchies. The pruning applied was three levels of root-children pruning, high p-value pruning with a threshold of 0.05, and zero-degree pruning.

The study reports that proteolysis and lipolysis are strongest in the Roquefort *P. roqueforti* population [31]. Proteolysis is reflected in alpha-amino acid (CHEBI:33704, *p_corr_* = 2.28 *×* 10*^−^*^30^), the most enriched class in the ‘Structure’ hierarchy, and in branched-chain amino acid (CHEBI:22918, *p_corr_* = 2.20 *×* 10*^−^*^10^), which comprises the precursors of the study’s proteolysis-derived aroma compounds, themselves reflected in methylbutanal (CHEBI:25282) and methylbutyric acid (CHEBI:38653, both *p_corr_* = 5.48 *×* 10*^−^*^3^) [31]. Lipolysis is reflected in the strong enrichment of fatty acid classes, such as saturated fatty acid (CHEBI:26607, *p_corr_* = 8.41 *×* 10*^−^*^21^). Lipolysis in blue cheeses produces aromatic compounds from the families of acids, methyl ketones, secondary alcohols and esters [31], which is reflected in the enrichment of methyl ketone (CHEBI:51867, *p_corr_* = 9.40 *×* 10*^−^*^12^), secondary alcohol (CHEBI:35681, *p_corr_* = 8.95 *×* 10*^−^*^4^) and several ester classes. The enriched classes from the ‘Role’ hierarchy, such as metabolite (CHEBI:25212, *p_corr_* = 1.44 *×* 10*^−^*^40^), are less specific but not surprising given the nature of the dataset. The high enrichment of plant metabolite (CHEBI:76924, *p_corr_* = 5.29*×*10*^−^*^19^) can be explained by the fact that many of the same metabolites detected in cheese also appear in plants, and thus carry plant-metabolite annotations in ChEBI. In contrast, crustacean metabolite and its two descendants *Daphnia* metabolite and *Daphnia magna* metabolite (CHEBI:83056, *p_corr_* = 9.31 *×* 10*^−^*^39^) are highly statistically significant despite being biologically spurious in this context. Checking the metabolites annotated with the role *Daphnia magna* metabolite reveals that these are among the most common metabolites found across living organisms, including several amino and fatty acids. That they carry this narrow, species-specific role in ChEBI rather than being annotated to every organism in which they occur, is a clear example of how incomplete and biased annotations in the ontology can lead to spurious enrichment. Overall, the enrichment graph provides a quick and representative overview of the major chemical classes highlighted by the blue-cheese study.

### Human dataset from Bernini et al. [32]

To analyse the use of the narrow backgrounds, we used a human-based dataset containing 14 metabolites from MTBLS46: *The Cardiovascular Risk of Healthy Individuals Studied by NMR Metabonomics of Plasma Samples* [32]. We examined it on both hierarchies using the full ChEBI background, the broad human background, and the more restrictive endogenous background; see Figures 4, S5, and 5 respectively. For the two narrow backgrounds we selected the option to expand the background to include all study-set leaves; otherwise, leaves outside the narrow background would have been excluded from the analysis. As a result, and since no pruning was applied, the same 130 structural classes were included in all three analyses; the number of role classes varied slightly, as role classes with no annotated leaf in the chosen background cannot be tested. Despite this, not all nodes are shown in all three graphs, since the default setting hides branches where all nodes are non-significant.

**Fig. 4.**
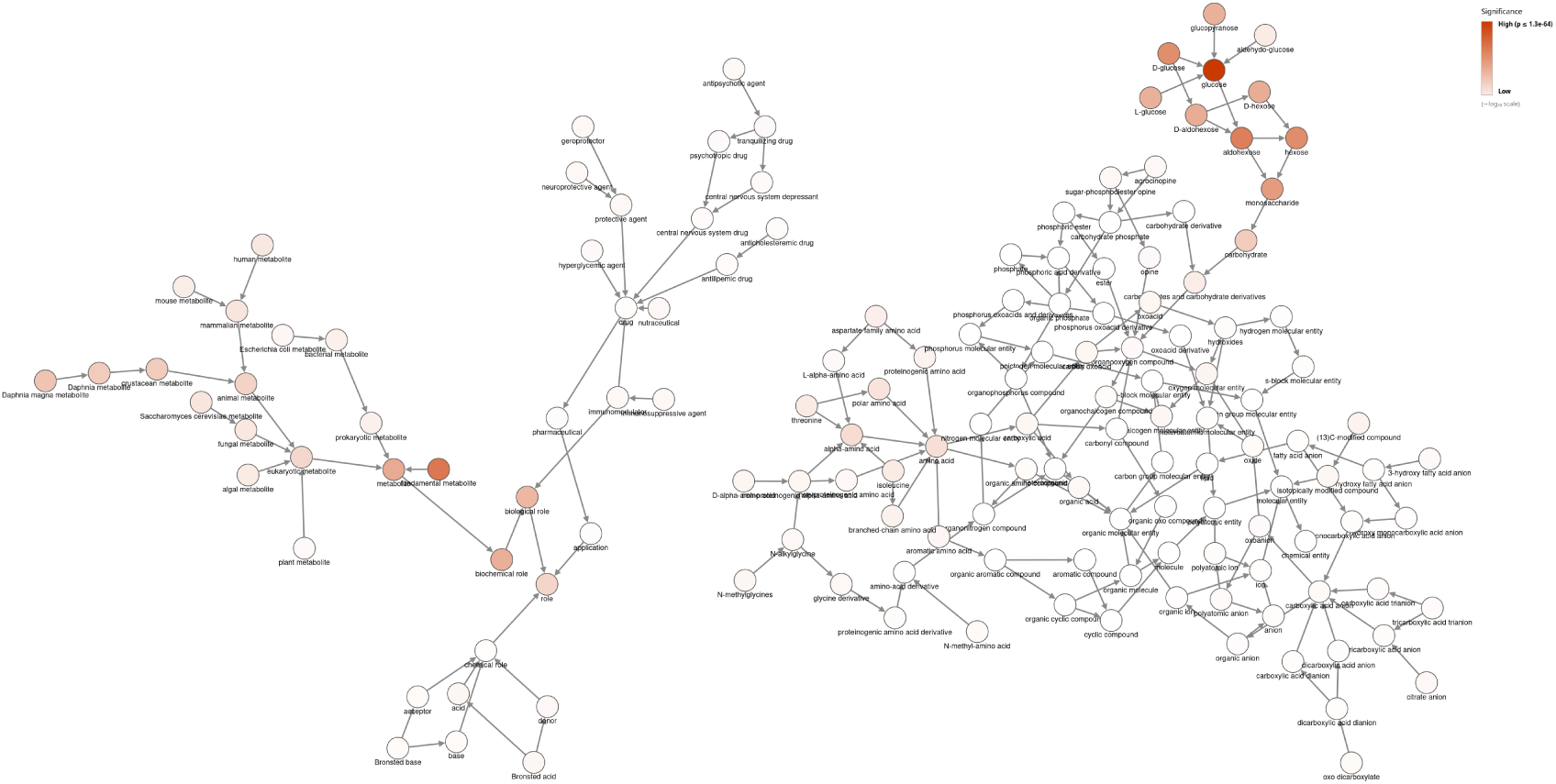
ChEBI-N enrichment analysis graph of the study-set from MTBLS46: *The Cardiovascular Risk of Healthy Individuals Studied by NMR Metabonomics of Plasma Samples* [32] on the full ChEBI background for both ‘Role’ and ‘Structure’ hierarchies. No pruning was applied.

**Fig. 5.**
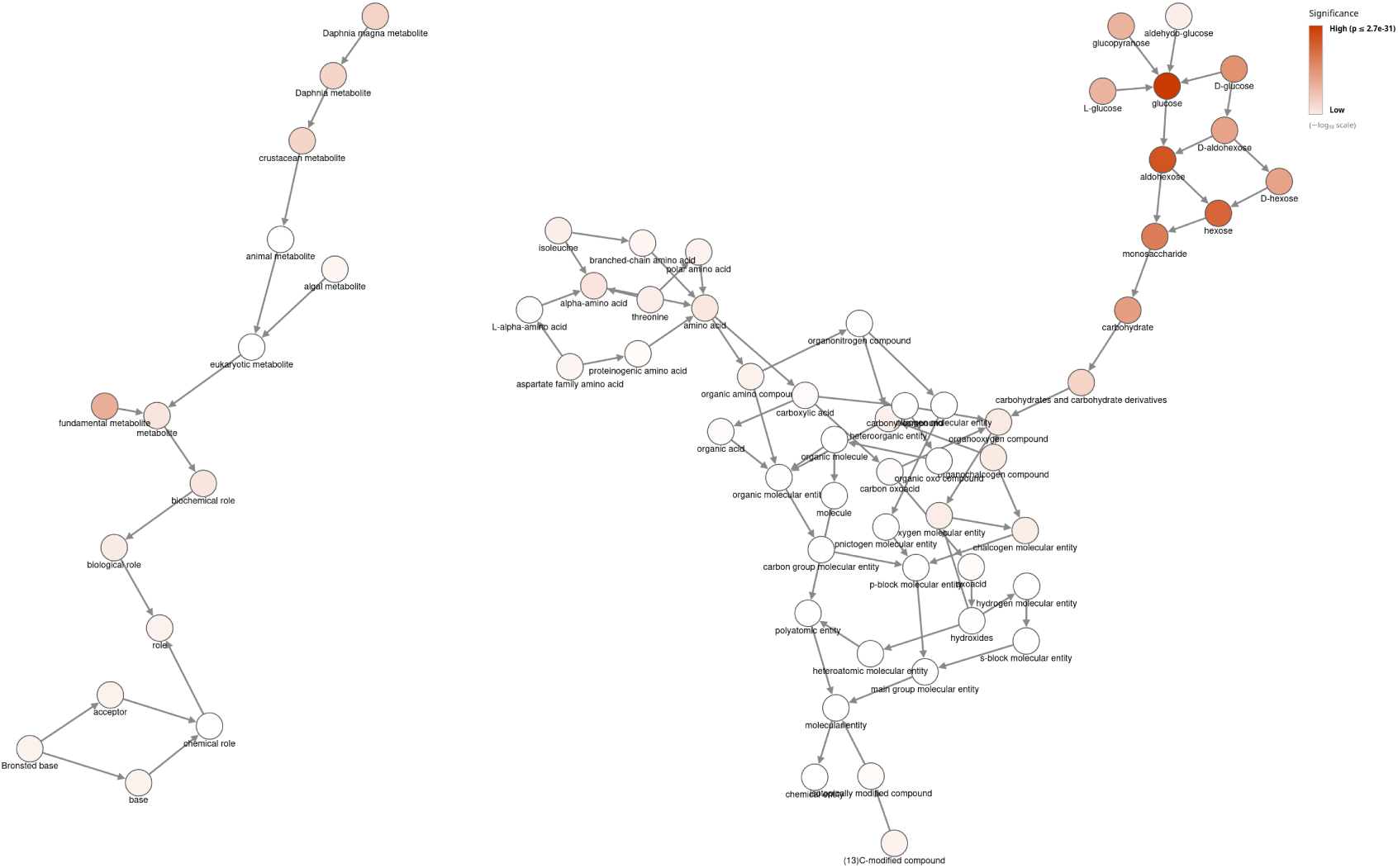
ChEBI-N enrichment analysis graph of the study-set from MTBLS46: *The Cardiovascular Risk of Healthy Individuals Studied by NMR Metabonomics of Plasma Samples* [32] on the narrow human background for both ‘Role’ and ‘Structure’ hierarchies. No pruning was applied.

Corrected p-values were generally lower, i.e. more enriched, for the largest background (full ChEBI) and higher, i.e. less enriched, for the narrowest one; this is apparent simply from the relative colour scales on each graph. These results are as expected since a smaller background size makes each entity proportionally less rare. There are far fewer significant classes (*p <* 0.05) for the endogenous background, with only 38 significant classes compared to 49 and 89 for the broader human and full ChEBI backgrounds, respectively. Any metabolite-like class is unremarkable against a human-based background, but appears enriched in the full ChEBI background. Examples of this include biological role (CHEBI:24432) for which corrected p-values of 1.0 *×* 10*^−^*^23^, 1.1 *×* 10*^−^*^7^, and 9.5 *×* 10*^−^*^4^ were obtained (in order of largest to smallest background), and human metabolite (CHEBI:77746) with corrected p-values of 5.5 *×* 10*^−^*^8^, 8.3 *×* 10*^−^*^1^, and 1.0. The same trend is visible for amino acid classes since the human backgrounds are already saturated with these, e.g. 5.3 *×* 10*^−^*^11^, 7.9 *×* 10*^−^*^5^, and 2.0 *×* 10*^−^*^4^ for amino acid (CHEBI:33709).

The glucose cluster is strongly enriched across all three backgrounds, with glucose (CHEBI:17234) being the most enriched class in all three cases, indicating that this result is robust. This aligns with the study’s finding that blood glucose (glycaemia) is one of the cardiovascular risk labels [32]. As for the cheese dataset, crustacean metabolite and its two descendants are highly statistically significant across all three backgrounds, again reflecting annotation biases rather than a biological signal.

Since the study [32] concerns human metabolites, the endogenous background may be the most fitting choice, whereas the full background inflates significance and, in effect, “rediscovers” that metabolites are metabolites. However, the differences in which classes are most enriched are not substantial, and the graphs look broadly similar, indicating that the full background is still a valid choice, particularly for studies on organisms that do not have a dedicated background available.

For both narrow backgrounds, most study-set leaves were absent and therefore added through background expansion. This is mainly because three study-set entries (glucose, threonine and isoleucine) are not leaf classes, so ChEBI-N replaces each of them with its leaf descendants, and many of those descendants are not present in the narrow backgrounds. Glucose alone has 17 leaf descendants. Only D-glucopyranose (CHEBI:4167) is present in both narrow backgrounds, and alpha-D-glucose (CHEBI:17925) and beta-D-glucose (CHEBI:15903) are present only in the broader human background. The added leaves make up less than 1% of each background, so the expansion does not change the backgrounds substantially.

### Limitations and future work

The results demonstrate the tool’s potential as a hypothesis-generating complement to metabolomics workflows, but the analyses also expose several limitations that point to directions for future development.

First, the enrichment results depend on the choice of background: the full ChEBI background tends to inflate the significance of generic classes. Improving how narrow backgrounds are built and validated is therefore a priority. In the future, users will be able to add their own backgrounds in the web interface, allowing results to be better tailored to the specific study. In addition, more pre-built organism-specific backgrounds will be added. We also intend to build a primary metabolism background from the metabolites shared by human, *Escherichia coli* and *Saccharomyces cerevisiae*. Since these metabolites can be expected to occur in most living organisms, such a background could complement those of organisms for which little species-specific metabolic data is available.

Second, the structural detail of the input sometimes does not match that of ChEBI. Most MS-based approaches identify metabolites only to the level of molecular connectivity, whereas ChEBI has separate classes for individual stereoisomers. Because matching is stereochemistry-sensitive, a connectivity-only SMILES matches either an unspecified leaf class or, via Chebifier, the leaves of its predicted parents; neither favours the more likely stereoisomer, which would be valuable to address in future work.

Finally, ChEBI-N’s results are only as reliable as the underlying ontology. As the analyses showed, incomplete or biased annotations can lead to spurious enrichments. The results should always be interpreted alongside domain expertise rather than in isolation. Moreover, ontology updates can change both the hierarchy and the set of annotated compounds, so p-values and the set of enriched classes may vary slightly between ChEBI releases. To support reproducibility, we recommend that analyses report the date of the most recent background update, as stated on the ChEBI-N webpage.

## 4 Conclusion

Like its predecessor BiNChE, ChEBI-N provides an interactive web tool for performing enrichment analysis on an input study-set based on the ChEBI ontology, and can also be used as a stand-alone Python library. This updated version additionally offers narrow backgrounds to adapt results to the study case, as well as the option to input SMILES or InChI as well as ChEBI IDs. ChEBI-N can assist in the interpretation of metabolomics study results, offering a visual graph that supports hypothesis formulation, and can serve as a complementary tool alongside existing metabolomics analysis pipelines. Future work includes expanding narrow background construction to further broaden its applicability to the metabolomics community.

## Supporting information

Supplementary Section

## Acknowledgements

We thank Simon Flügel (University of Osnabrü ck) for help with integrating Chebifier into ChEBI-N. This study was funded by the SNF under grant agreement MetaboLinkAI (grant agreement number 10002786).

## Notes

### Competing Interest Statement

The authors have declared no competing interest.

