## Supplementary Section for "ChEBI-N - Enrichment Analysis with the ChEBI Ontology"

#### S1 Ontology visualisation

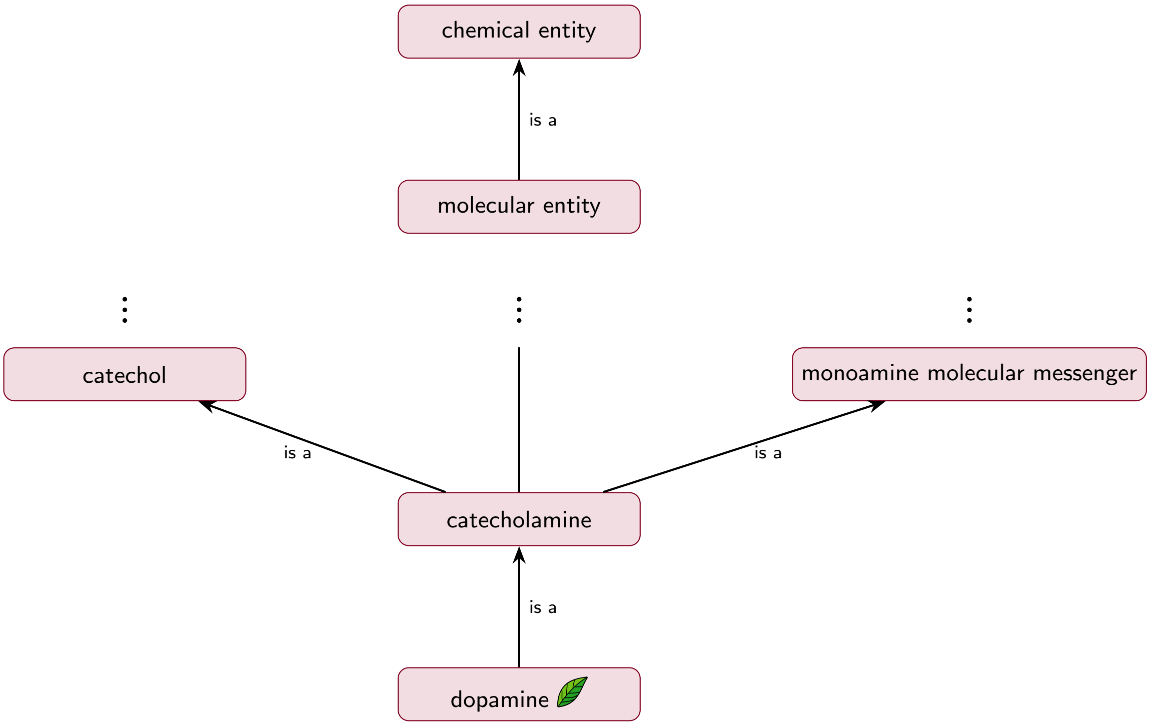


**Fig. S1** A visual depiction of an excerpt from the chemical entity hierarchy in the ChEBI ontology. Each class is linked to its parent class(es) through *is a* relationships. The dots indicate levels of the hierarchy not shown. Note that although the arrows point upwards, towards the root, we refer to a class as a descendant (or child) of the classes above it.

#### S2 Leaf classes in ChEBI-N background

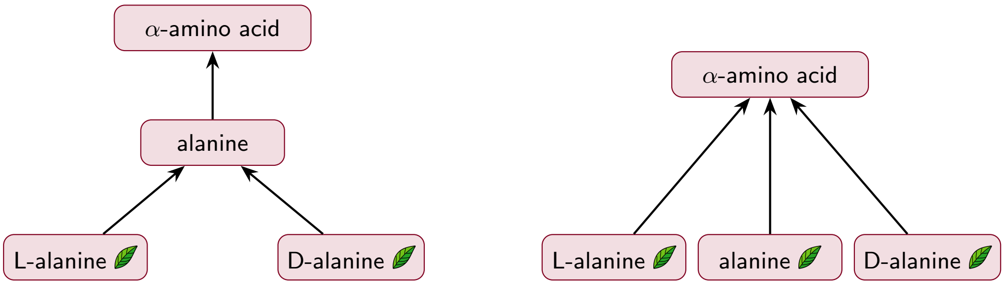


(a) Original ChEBI hierarchy (b) Hierarchy used for the ChEBI-N background

**Fig. S2** Flattening of structurally defined classes, illustrated with alanine. Leaf classes are marked with a leaf symbol. (a) In the original hierarchy, alanine is a parent of L-alanine and D-alanine, so only the latter two are leaves. (b) In the hierarchy used for the background, the edges from L-alanine and D-alanine to alanine are replaced by edges to the parent of alanine, so that all three structurally defined classes become sibling leaves.

#### S3 ChEBI-N graph screenshot

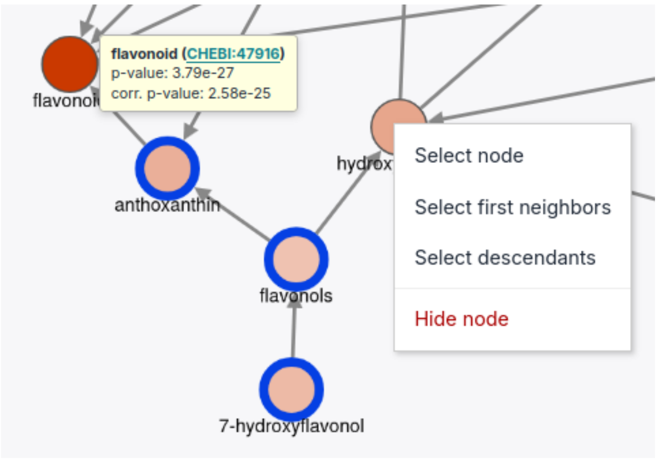


**Fig. S3** Zoomed-in screenshot of a results graph from ChEBI-N. The flavonoid node has the mouse hovered over it, the node to the right has been right-clicked, and the nodes framed in blue have been selected by left-clicks.

#### S4 p-value calculations

To calculate p-values for non-weighted study-sets, we use Fisher's exact test, which answers the question: which categories are more common among the molecules in the study-set than would be expected by a chance sample of the given background distribution? We compute the counts in the contingency table, as shown in Table S1, for each class in question and use them in the calculations. We perform calculations for the different hierarchies ('Structure' and 'Role') separately.

**Table S1** Contingency table for enrichment of a class A. An observation cannot fall into more than one category represented in the table. Leaf classes are referred to as leaves.

| | Leaves in class A | Leaves not in class A | Total |
|---|---|---|---|
| study-set | $a$ | $b$ | $a+b$ |
| Background | $c$ | $d$ | $c+d$ |
| Total | $a+c$ | $b+d$ | $n$ |

$P$ is the exact probability of the observed table, and is calculated as follows using the values from the table:

$$P = \frac{(a+b)!\,(c+d)!\,(a+c)!\,(b+d)!}{a!\,b!\,c!\,d!\,n!} \tag{S1}$$

For weighted calculations, the SaddleSum method [1], which implements the Lugannani-Rice formula, is used to compute enrichment p-values. We translated and adapted the C code of Stojmirović and Yu [1]. SaddleSum's p-value for a class depends on which of the weighted study-set members fall under it and how heavily weighted they are, not on how large or specific the class itself is. For detailed explanations of the mathematical foundations, see Stojmirović and Yu [1].

#### S5 Pruning strategies

After inserting the study-set, the user can choose if they want to use any pruning methods to make the final graph less cluttered. Some of the pruning methods that are not based on p-values will be applied before any p-value calculations are made to reduce computation. Any, none or all of the individual pruning methods can be selected.

The pruning methods that are applied before p-value calculations are the following.

**Root Children Pruner:** Removes the root(s) and their descendants up to the chosen level; for example, choosing three levels removes the roots and two levels of their descendants. This allows the removal of more general, and thus less meaningful, entities in the ontology.

**Linear Branch Collapser Pruner:** Removes linear branches within the graph, meaning that only nodes with one directly ascending (parent) node and one directly descending (child) node will be removed. Every $n$th intermediate node in the branch will be kept, where $n$ can be set to zero. Setting $n = 0$ means that only the first and last nodes will be kept.

After p-values are calculated for the nodes that have not been removed, the following pruners are applied, if chosen.

**High p-Value Branch Pruner:** Removes branches from the graph that only contain nodes with a p-value greater than the threshold (the standard threshold being 0.05, but this can be changed). A node with a higher p-value will thus be kept if it has at least one descendant with a p-value lower than the threshold.

**Zero-degree Pruner:** Removes nodes that have no connections with other nodes; i.e. nodes with no connecting neighbours.

If pruning methods are manually selected, they are used once each, in the following order: root-children → linear-branch → (enrichment-calculations + p-value correction) → high-p-value → zero-degree. Only the selected pruners will be applied. Alternatively, they can be applied in a predefined looping manner:

**Plain Enrichment Pruning Strategy:** First there is a pre-loop phase, where pruners are applied once, and then a loop phase, where pruners are applied repeatedly until no more changes are made. The pre-loop phase applies the high p-value branch pruner with a threshold of 0.05, the linear branch collapser pruner (n = 0), and the root children pruner (levels = 2). The loop phase applies the high p-value branch pruner with a threshold of 0.05, the linear branch collapser pruner (n = 0), and the zero-degree pruner. p-values are calculated for all classes before the pre-loop phase. At the start of each loop iteration, Benjamini-Hochberg correction is recomputed over the classes that remain, and the high p-value pruner uses these updated corrected p-values.

When the graph is displayed, the user can choose to re-run the calculations with new pruning options under *Settings*. In addition, an interactive p-value filter in the form of a slider is available in the graph view to easily adapt which nodes are visible based on their significance.

### S6 Additional ChEBI-N result graphs

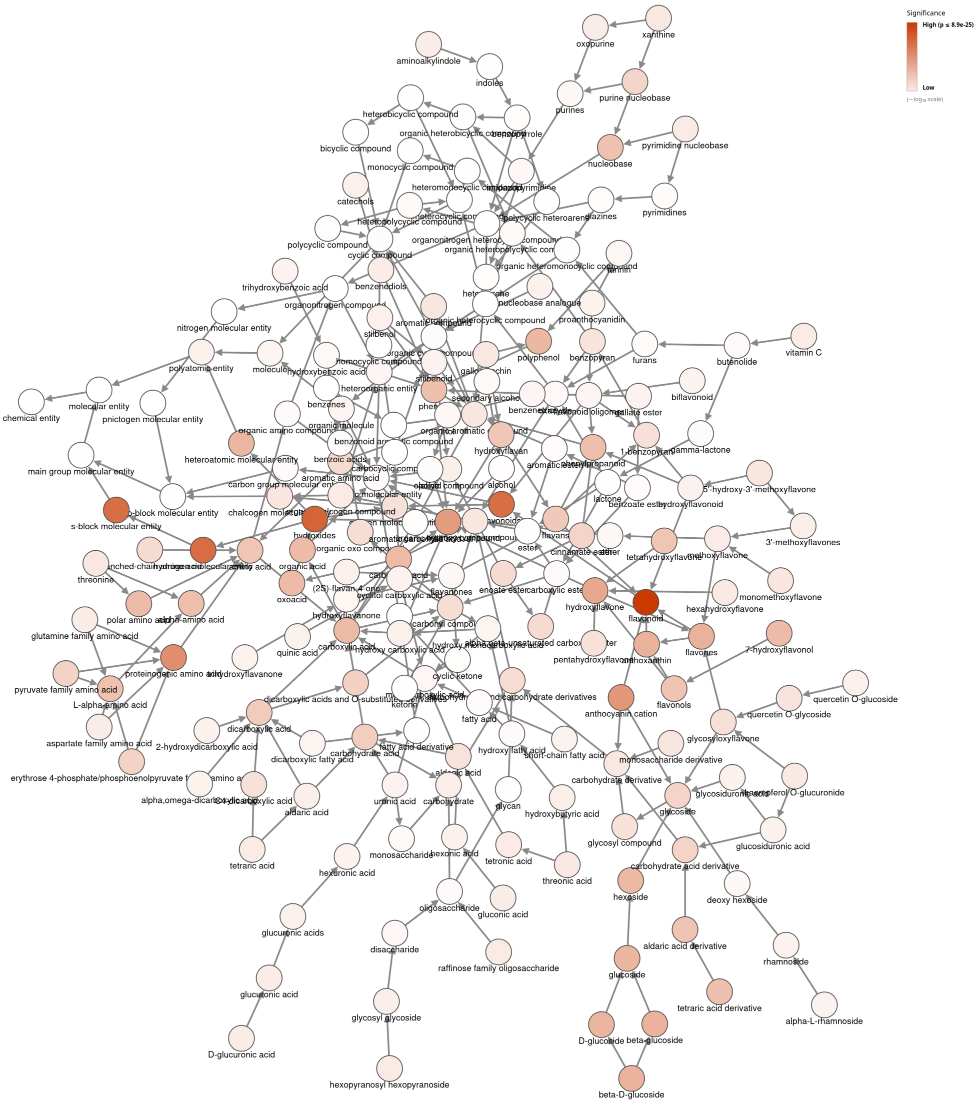


**Fig. S4** ChEBI-N enrichment analysis graph of the study-set from MTBLS55: *The influence of storage on the "chemical age" of red wines* [2] using ChEBI IDs. The results are only on the 'Structure' hierarchy, and no pruning strategies have been applied.

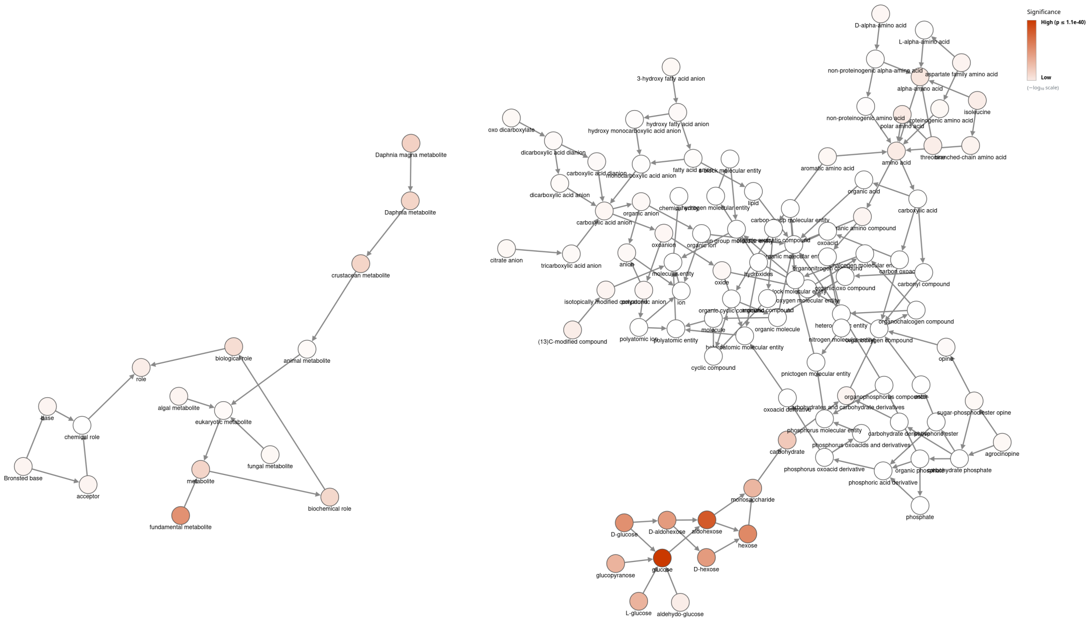


**Fig. S5** ChEBI-N enrichment analysis graph of the study-set from MTBLS46: *The Cardiovascular Risk of Healthy Individuals Studied by NMR Metabonomics of Plasma Samples* [3] on the broad human-based background for both 'Role' and 'Structure' hierarchies. No pruning was applied.

### S7 Applying ChEBI-N on a larger dataset

##### *Human dataset from Shcherbakov et al. [4]*

To evaluate ChEBI-N's performance on a substantially larger study-set, we applied it to data from MTBLS106, the metabolomics dataset associated with Shcherbakov et al.'s study on ribosomal mistranslation [4]. Of the 586 metabolites reported in the study, 564 had annotated ChEBI identifiers and were used for analysis. Given the considerably larger study-set size relative to the other datasets tested, we anticipated an increase in the number of significantly enriched classes and correspondingly greater visual density in the resulting enrichment graph.

We first performed enrichment on both the 'Role' and 'Structure' hierarchies against the full ChEBI background, applying the plain enrichment pruning strategy; see Figure S6.

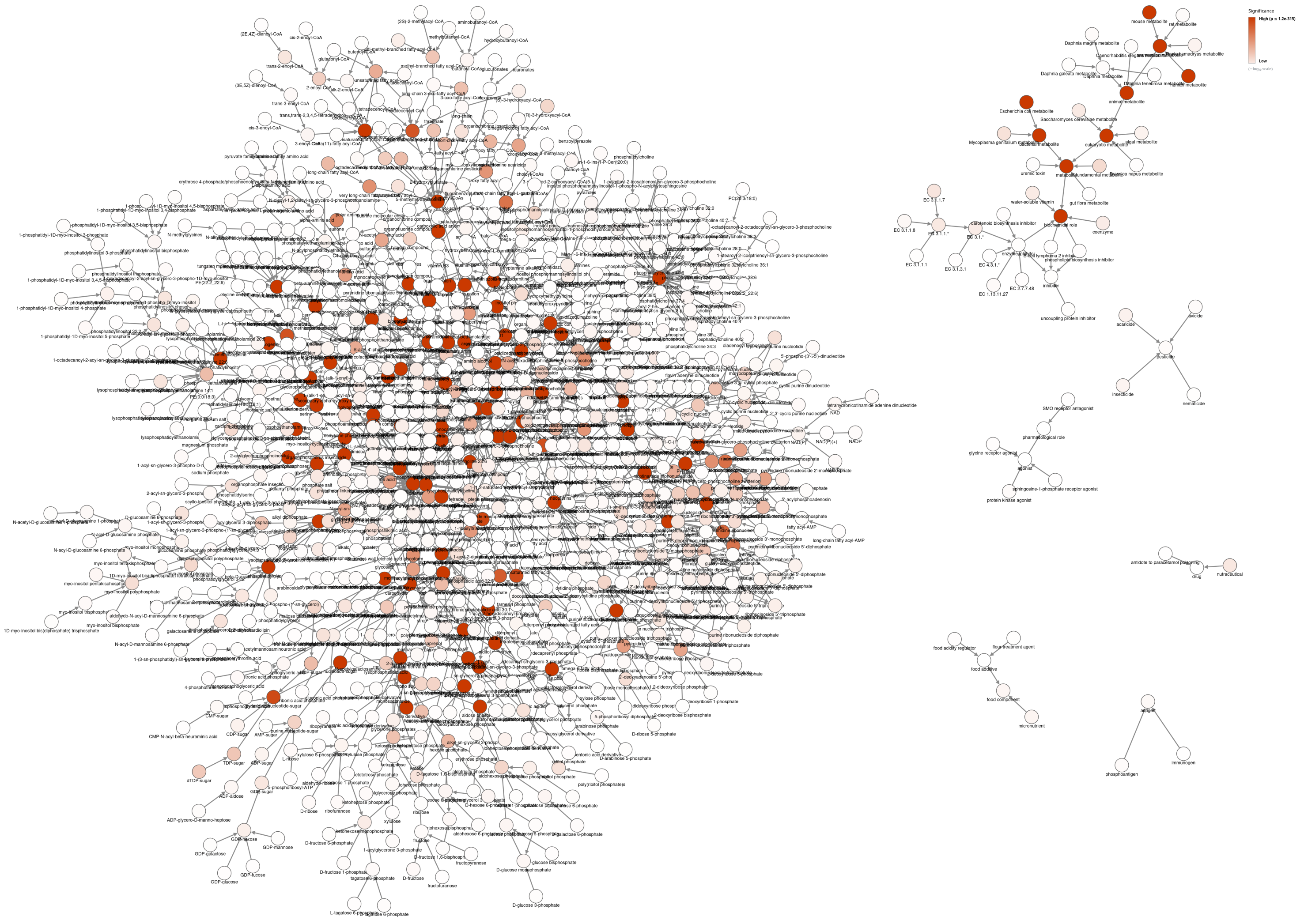

**Fig. S6** Enrichment analysis of the study-set from MTBLS106: *Ribosomal mistranslation leads to silencing of the unfolded protein response and increased mitochondrial biogenesis* [4] on both ‘Role’ and ‘Structure’ hierarchies on the full background. The plain enrichment pruning strategy (looping pruning method) was applied.

As anticipated, the resulting graph was densely populated, making individual classes difficult to distinguish. By default, all nodes with $p \leq 0.05$ are retained, together with their ancestor paths to the root; for a result set of this magnitude where the scale extends down to $1.2 \times 10^{-315}$ this default threshold gives limited discriminatory value. To address this, we used the interactive p-value slider to progressively filter out nodes from display. Figure S7 shows the same analysis filtered to $p \leq 4.0 \times 10^{-131}$ (not keeping paths to root). While still visually dense, this filtered view was considerably more manageable.

We then repeated the analysis using the narrow human background (with background expansion), applying the same pruning method; see Figure S8, filtered to hide nodes with $p > 1.2 \times 10^{-37}$. Consistent with the comparison of background choice on the human dataset by Bernini et al. [3], as discussed in the Results and discussion section of the main text, this analysis also shows that many classes were less significant under the narrower background, reducing overall noise in the graph. Comparisons of the graphs show that substantially more nodes have been pruned under the narrow background, indicating that a considerable number of classes lose significance once the background is restricted, and the resulting graph is thus easier to interpret.

Worth noting, however, is that several classes received p-values of 0.0000e+00. As an actual p-value of zero is highly unlikely, this instead indicates that the true value is too small to be represented in floating-point precision. This occurs primarily for generic classes positioned high in the hierarchy, such as `pnictogen molecular entity` (CHEBI:33302), `phosphorus molecular entity` (CHEBI:26082), and `oxygen molecular entity` (CHEBI:25806). This makes it more difficult to identify which of the more specific, biologically relevant classes are the strongest hits for this particular study.

A related consequence of this large p-value range is that the colour scale adapts accordingly. Without this adaptation, an even larger number of classes would appear fully red and be visually indistinguishable from one another. However, this adaptation introduces its own trade-off: classes with genuinely very small p-values can now appear only faintly coloured, simply because the scale has been stretched to accommodate the more extreme values.

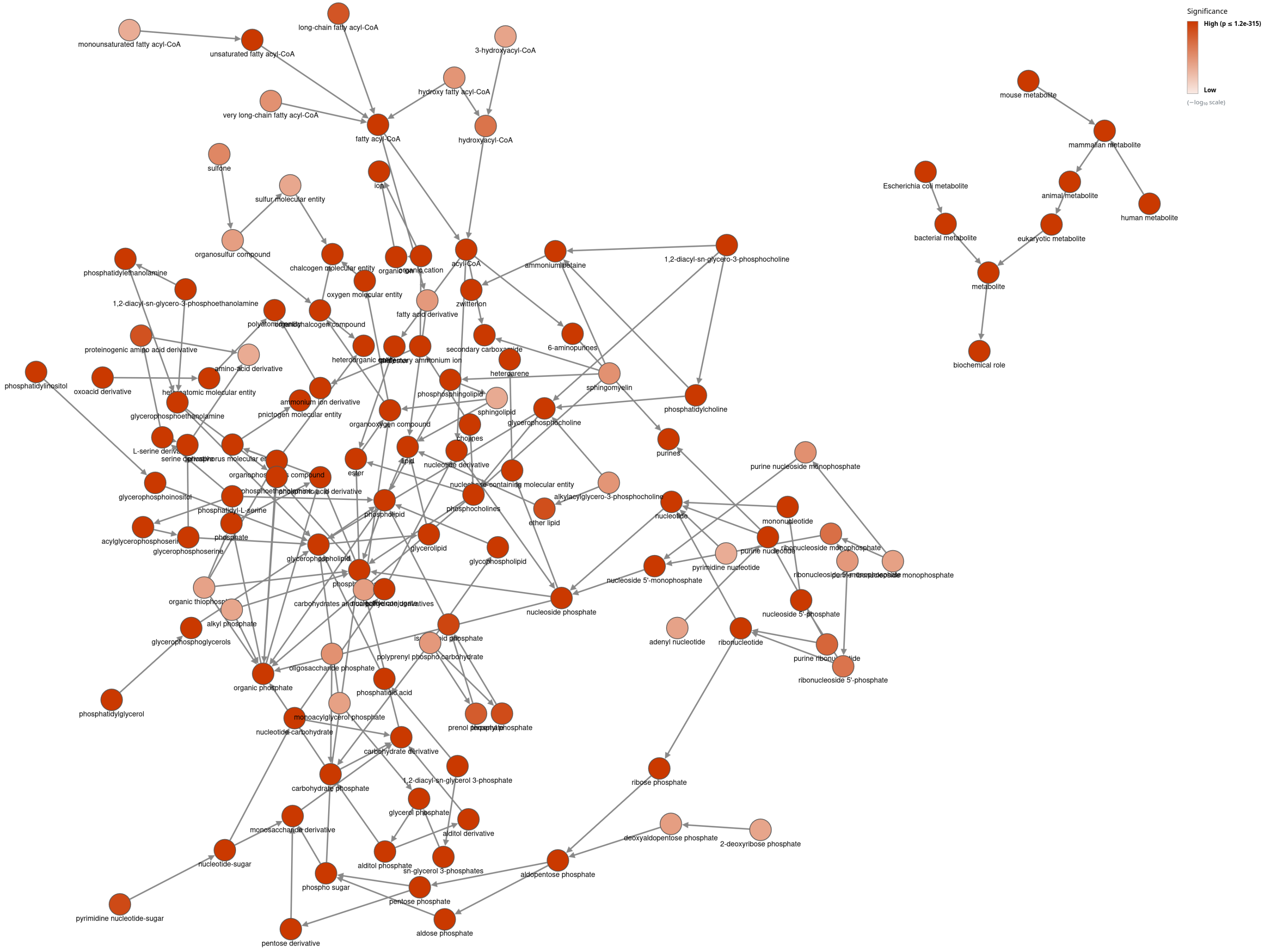


**Fig. S7** Enrichment analysis of the study-set from MTBLS106: *Ribosomal mistranslation leads to silencing of the unfolded protein response and increased mitochondrial biogenesis* [4] on both 'Role' and 'Structure' hierarchies on the full background. The plain enrichment pruning strategy (looping pruning method) was applied and the dynamic p-value filter was used, hiding nodes with p-values greater than $4.0 \times 10^{-131}$.

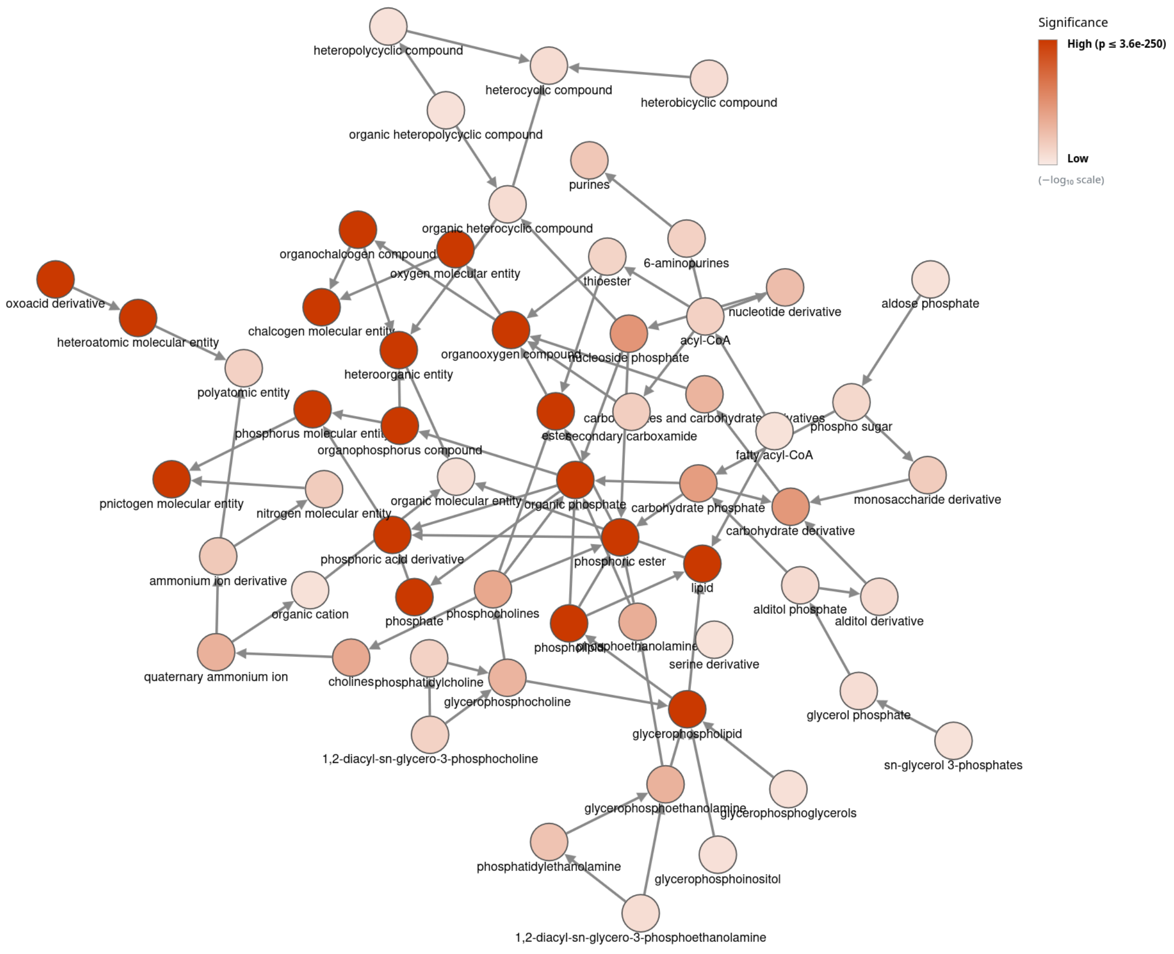


**Fig. S8** Enrichment analysis of the study-set from MTBLS106: *Ribosomal mistranslation leads to silencing of the unfolded protein response and increased mitochondrial biogenesis* [4] on both 'Role' and 'Structure' hierarchies on the narrow human background. The plain enrichment pruning strategy (looping pruning method) was applied and the dynamic p-value filter was used, hiding nodes with p-values greater than $1.2 \times 10^{-37}$.
